# Antimicrobial resistance, virulence genes profiles and molecular epidemiology of carbapenem-resistant *Klebsiella pneumoniae* strains from captive giant pandas (*Ailuropoda melanoleuca*)

**DOI:** 10.1101/2024.02.20.581254

**Authors:** Xia Yan, Mei Yang, James Edward Ayala, Lin Li, Yang Zhou, Rong Hou, Songrui Liu, Yunli Li, Chanjuan Yue, Dongsheng Zhang, Xiaoyan Su

## Abstract

This study aimed to investigate the antibiotic susceptibility, antibiotic resistance genes (ARGs), mobile genetic elements (MGEs), virulence genes, and molecular epidemiology of carbapenem-resistant *Klebsiella pneumoniae* (CRKP) strains isolated from giant pandas. The screening of 178 nonduplicated *Klebsiella pneumoniae* strains identified eight CRKP strains, with the most abundant ARGs observed in ampC/blaDHA, blaSHV-01, blaSHV-02, tetB-01, tetB-02, tetC-01, and tetC-02. MGE analysis revealed the presence of intI1 in all strains, while other MGEs exhibited varying detection rates. Strain 24 exhibited the highest diversity in terms of MGE species. Seven virulence genes including wabG, uge, ycf, entB, kpn, alls, and wcaG, showed positive results with different proportions across the strains. Molecular epidemiology analysis using pulsed-field gel electrophoresis (PFGE) patterns indicated a high level of genetic diversity among the CRKP strains. Multi-locus sequence typing (MLST) analysis classified the strains into different sequence types (STs). In conclusion, this study highlighted the diverse nature of CRKP strains found in giant pandas, which exhibited varying levels of antibiotic resistance along with multiple ARGs and virulence genes present. These findings emphasized the importance of monitoring and researching antibiotic resistance within wildlife populations to safeguard the health status of these endangered animals.

## 1. Introduction

The giant panda, an endemic species to China, serves as the flagship species for global wildlife conservation efforts (Jia W et al. 2023). Bacterial diseases have emerged as a major epidemic that poses a grave threat to the life and well-being of giant pandas. With the widespread utilization of antibiotics, bacterial resistance continues to emerge and escalate every year, thereby posing an immense risk to the health of giant panda (Yang X et al. 2017; Guo L et al. 2015). Gastrointestinal bacterial infections are believed to be a primary cause of mortality among giant pandas, with alterations in gut microecology leading to digestive disorders and potential disease development. The intestinal disease in giant pandas can be caused by bacterial pathogens such as *Escherichia coli* (*E. coli*), *Klebsiella pneumoniae* (KP), *Campylobacter jejuni, Arizona bacillus, Pseudomonas aeruginosa* and other bacteria (Xiong Yan et al. 2000).

KP is a gram-negative, conditionally pathogenic bacillus. In captive giant pandas, KP infection has become increasingly prevalent and often occurs in conjunction with other bacteria, making it the most important pathogen (Wang Qiang et al. 1998; Wang Chengdong et al. 2006; Wang Xiaoyu et al. 2002). Infected pandas may develop haemorrhagic enteritis characterized by bloody stools and genitourinary bleeding characterized by haematuria, which can potentially result in fatal haemorrhagic sepsis. In addition, our previous study has identified the emergence of drug resistance in giant panda-derived KP (Xiong Yan et al. 1998).

According to our research, antimicrobials have been extensively utilized for the prevention and treatment of infectious diseases in giant pandas over the past few decades (Yang X et al. 2017; Guo L et al. 2015). The misuse of antibiotics is the primary factor contributing to the emergence of carbapenem-resistant *Klebsiella pneumoniae* (CRKP). Furthermore, horizontal gene transfer (HGT) via mobile genetic elements (MGEs) such as integrons, transposons, integration-coupled elements, genomic islands and plasmids plays an important role in disseminating antibiotic resistance genes (ARGs) carried by MDR-KP (Chen Q et al. 2016; Partridge et al. 2009). Integrons possess the ability to capture, transform and adapt one or more resistance gene cassettes into functionally expressed genes through self-acting gene expression systems (Gillings et al. 2014). Their association with plasmids also facilitates the transfer of these genes among different bacterial species. The three main types of MGEs associated with antimicrobial resistance are classified as type 1, type 2 and type 3 integrons (Kaushik et al. 2018).

Recent studies have identified a large number of antimicrobial resistant bacteria (ARB), ARGs and their MGEs (including integrons) in *E. coli* isolated from captive giant pandas (Yang, X. et al. 2016; Yang, X. et al. 2018; Mustafa et al. 2021). However, limited information is available regarding the prevalence of CRKPs, the diversity of ARGs and MGEs, and the correlation between antimicrobial resistance and the occurrence of integron gene cassettes in CRKP among captive giant pandas (Yang, X. et al. 2016). Additionally, there is a lack of knowledge about the antimicrobial resistance profiles across different age groups of giant pandas. The objective of this study was to analyze the antimicrobial resistance profiles of 178 KP strains collected from fecal samples obtained from captive giant pandas belonging to various age and sex groups. Furthermore, we aimed to investigate the presence of ARGs, integrative subgene cassettes and other MGEs in 8 CRKP strains. These findings will provide valuable insights for guiding appropriate use of clinical antibiotics in giant pandas.

## 2. Materials and Methods

### 2.1 Bacterial isolates and screening for carbapenemases phenotype

One hundred seventy-eight nonduplicated KP isolates were collected from fresh feces of captive giant pandas at the Chengdu Research Base of Giant Panda Breeding (Panda Base) in Sichuan, China, between 2018 and 2019. These isolates were identified as KP by Gram staining, 16S rDNA analysis and bacterial biochemical identification. Carbapenemases production was screened in all isolates using cefotaxime (CTX) and ceftazidime (CAZ) alone according to CLSI guidelines (2019). The presence of carbapenemases in the isolates was determined phenotypically by observing diameter enhancement of the inhibition zone around the clavulanate disk and corresponding β-lactam antimicrobial disk. If the enhancement value exceeded 5 mm, the isolate was considered an carbapenemases producer.

### 2.2 Antimicrobial susceptibility testing of CRKP isolates

The antibiotic resistance testing of CRKP isolates was performed using the disk diffusion method (K-B method) against a panel of antibiotics, including piperacillin (PIP), moxalactam (MOX), ceftazidime (CAZ), cefixime (CFM), ceftazidime (CMZ), cefepime (FEP), cefotaxime (CTX), cephalexin (CA), cephazolin (CZ), ceftriaxone (CTR), cefoxitin (FOX), piperacillin/tazobactam (TZD), cefuroxime (CXM), cefaclor (CEC), ampicillin/sulbactam (AMS), cefoperazone (CFP), ceftizoxime (ZOX), aztreonam (AT), meropenem (MEM), imipenem (IPM), kanamycin (K), gentamicin (GM), streptomycin (S), enoxacin (ENX), ofloxacin (OFX), norfloxacin (NOR), lomefloxacin (LOM), fleroxacin (FO), levofloxacin (LVX), ciprofloxacin (CIP), gatifloxacin (GAT), chloramphenicol (C), azithromycin (AZM), doxycycline (DX), minocycline (MI), compound sulfamethoxazole (SXT), trimethoprim (TMP). The antibiotic disks were purchased from Hangzhou Microbiological Reagent Co. Ltd, Hangzhou, China. *E. coli* ATCC25922 was used as the quality control bacterial strain. The results we interpreted as susceptible (S), intermediate (I), and resistant (R) based on the interpretative criteria of the CLSI 2020.

### 2.3 DNA extraction of CRKP isolates

The CRKP isolates were subjected to total genomic DNA extraction using the TIANamp Bacteria DNA Kit (Tiangen Biotech, Beijing, China) following the manufacturer’s protocol. Subsequently, the DNA samples were stored at -20 °C.

### 2.4 Antibiotic resistance genes analysis of CRKP isolates

High-throughput qPCR (HT-qPCR) reactions were conducted using the Wafergen smartchip Real-time PCR system to analyze the antibiotic resistance genes. A total of 89 primer sets were employed, which were listed in supplementary Table 1 for the detection of resistance genes. Each sample was simultaneously replicated three times. Following a pre-denaturation step at 95 °C for 10 min, amplification was performed through 30 cycles according to the following program: denaturation at 95 °C for 30 s, annealing at 60 °C for 30 s. The obtained results were then analyzed with smartchip qPCR Software to exclude wells exhibiting multiple melting peaks or amplification efficiency beyond the range (90%-110%).

### 2.5 The Mobile genetic elements (MGEs) analysis of CRKP isolates

Twenty-three pairs of primers were used to detect MGEs in CRKP isolates by PCR, following previously described protocols (Levesque, Piche et al. 1995, White, McIver et al. 2001, Zou, Li et al. 2018, Zhu, Pan et al. 2020). The sequences of primers were listed in supplementary Table 1. PCR amplification was performed in a total volume of 25 μL containing 12.5 μL of Dream Taq Green PCR Master Mix (2×), 8.5 μL ddH_2_O, and each forward primer and reverse primer at a concentration of 1 μL, DNA template 2 µL. Amplification was carried out under the following thermal cycling conditions: pre-denaturation at 95 °C for 5 min, followed by a total of 30 cycles consisting of denaturation at 95 °C for 30 s, annealing at the specified temperature for 30 s, extension at 72 °C for 30 s, and final extension at 72 °C for 10 min. PCR products were subsequently subjected to 1% agarose gel at 120 V, 0.5 × TAE buffer electrophoresis for 38 min.

### 2.6 The virulence gene analysis of CRKP isolates

The multiplex PCR reaction mixtures for 16 virulence genes (magA-fimH-uge-iutA, wabG-rmpA-cnf-ycf, hly-iroN-K2a-mrkD, kpn-allS-entB-wcaG) in CRKP isolates were divided into four different sets (Candan ED et al. 2015). Each set consisted of a total volume of 50 μL containing 25 μL of Dream Taq Green PCR Master Mix (2×), 2 μL of total DNA, 22 μL ddH_2_O, and each forward primers and reverse primers (Sangon Biotech Co., Ltd., Shanghai, China) at a concentration of 0.5 μL. The amplification process was carried out with the following thermal cycling conditions: pre-denaturation at 95 °C for 5 min, followed by 30 cycles consisting of denaturation at 94 °C for 1 min, annealing at 58 °C for 1 min, extension at 72 °C for 1 min, and final elongation at 72 °C for 10 min. Finally, the PCR products were subjected to 2% agarose gel at 120 V, 1 × TAE buffer electrophoresis for 38 min.

### 2.7 The molecular epidemiology analysis of CRKP isolates

PFGE was conducted to investigate the molecular epidemiology of the CRKP isolates with XbaI-digested and genotyped DNA. The genomic DNA restriction patterns of the isolates were analyzed and interpreted based on the previously established criteria (Han, Zhou et al. 2013). Additionally, in order to further assess whether clonal spread influenced the dissemination of carbapenemase-producing KP isolates in giant panda, MLST was performed by amplifying internal fragments of seven K. pneumoniae housekeeping genes provided on the MLST website (http://www.mlst.net).

## 3. Result

### 3.1 Antibiotic susceptibility of CRKP isolates

The screening and detection process yielded a total of 8 CRKP isolates, with a isolation rate 4.5% (8/178). Prior to further analysis, we conducted an initial assessment of antibiotic susceptibility. The results of antibiotic susceptibility revealed that seven of eight CRKP strains (strain 24, 74, 77, X25, X41, X46, E28) exhibited resistance to imipenem, while one strain (strain 45) showed resistance to meropenem. Additionally, strain 24 demonstrated multiple resistance with a spectrum including imipenem, chloramphenicol, doxycycline, minocycline, compound sulfamethoxazole, and trimethoprim. Strain E28 displayed widely resistance to β-lactamase such as cephalexin, cefazolin, cefoxitin, and imipenem. In addition to meropenem, strain 45 was resistant to ceftazidime and doxycycline. However, strain 74, 77, X25, X41, X46 didn’t exhibit any resistance except for imipenem (Fig. 1).

**Fig. 1.**
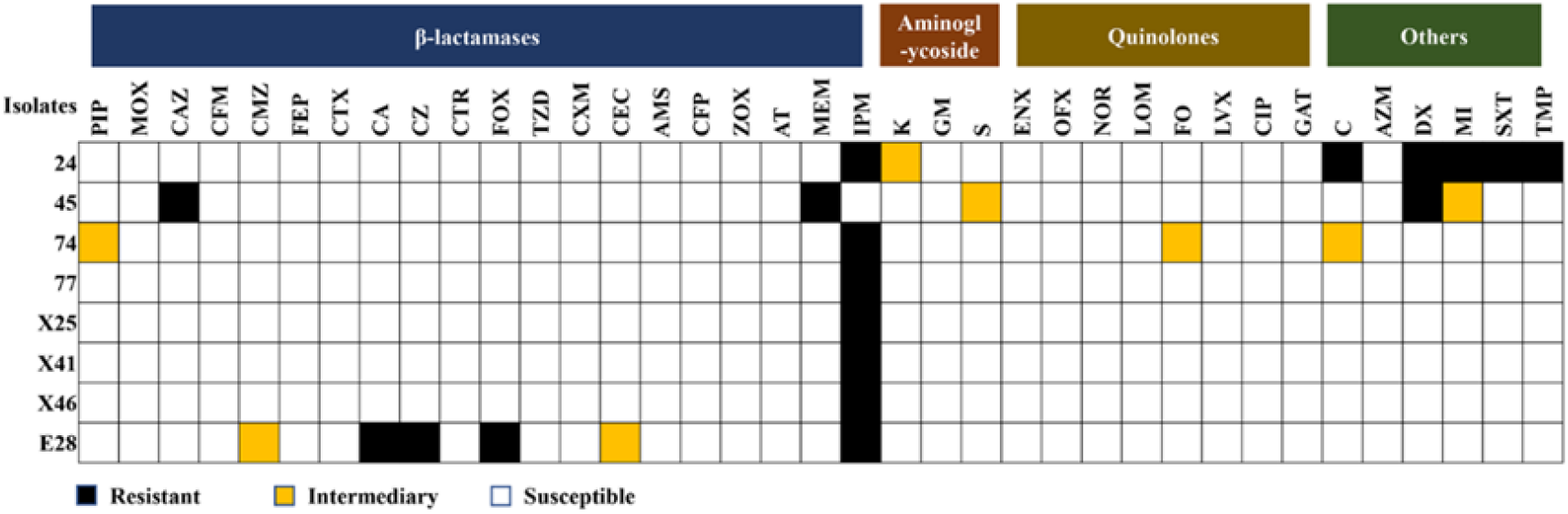
The antibiotic susceptibility profiles of CRKP isolates. Columns: 37 different types of antibiotics; Rows: the number of isolates in the study. PIP: piperacillin, MOX: moxalactam, CAZ: ceftazidime, CFM: cefixime, CMZ: ceftazidime, FEP: cefepime, CTX: cefotaxime, CA: cephalexin, CZ: cephazolin, CTR: ceftriaxone, FOX: cefoxitin, TZD: piperacillin/tazobactam, CXM: cefuroxime, CEC: cefaclor, AMS: ampicillin/sulbactam, CFP: cefoperazone, ZOX: ceftizoxime, AT: aztreonam, MEM: meropenem, IPM: imipenem, K: kanamycin, GM: gentamicin, S: streptomycin, ENX: enoxacin, OFX: ofloxacin, NOR: norfloxacin, LOM: lomefloxacin, FO: fleroxacin, LVX: levofloxacin, CIP: ciprofloxacin, GAT: gatifloxacin, C: chloramphenicol, AZM: azithromycin, DX: doxycycline, MI: minocycline, SXT: compound sulfamethoxazole, TMP: trimethoprim.

### 3.2 Antibiotic resistance genes in 8 CRKP isolates

A total of 89 ARGs were assessed in 8 CRKP isolates using HT-qPCR. Out of these, 47 ARGs were detected, 6 ARGs among which were positively present in all strains, including aacC, blaCTX-M-04, blaOXY, blaSHV-01, blaSHV-02 and vanTC-02. However, 43 ARGs such as aac (6’)-II, aac (6’)-Iy, aacC1, blaPER were not detected. Furthermore, the abundance of the identified 47 positive ARGs varied among the strains. The top ten resistance genes of abundance were: aacC, ampC/blaDHA, blaSHV-01, blaSHV-02, tetB-01, tetB-02, tetC-01, tetC-02, tetD-02, and vanTC-02. It was worth noting that ARGs of tetracycline exhibited highly abundance in strain 45 (Fig. 2).

**Fig. 2.**
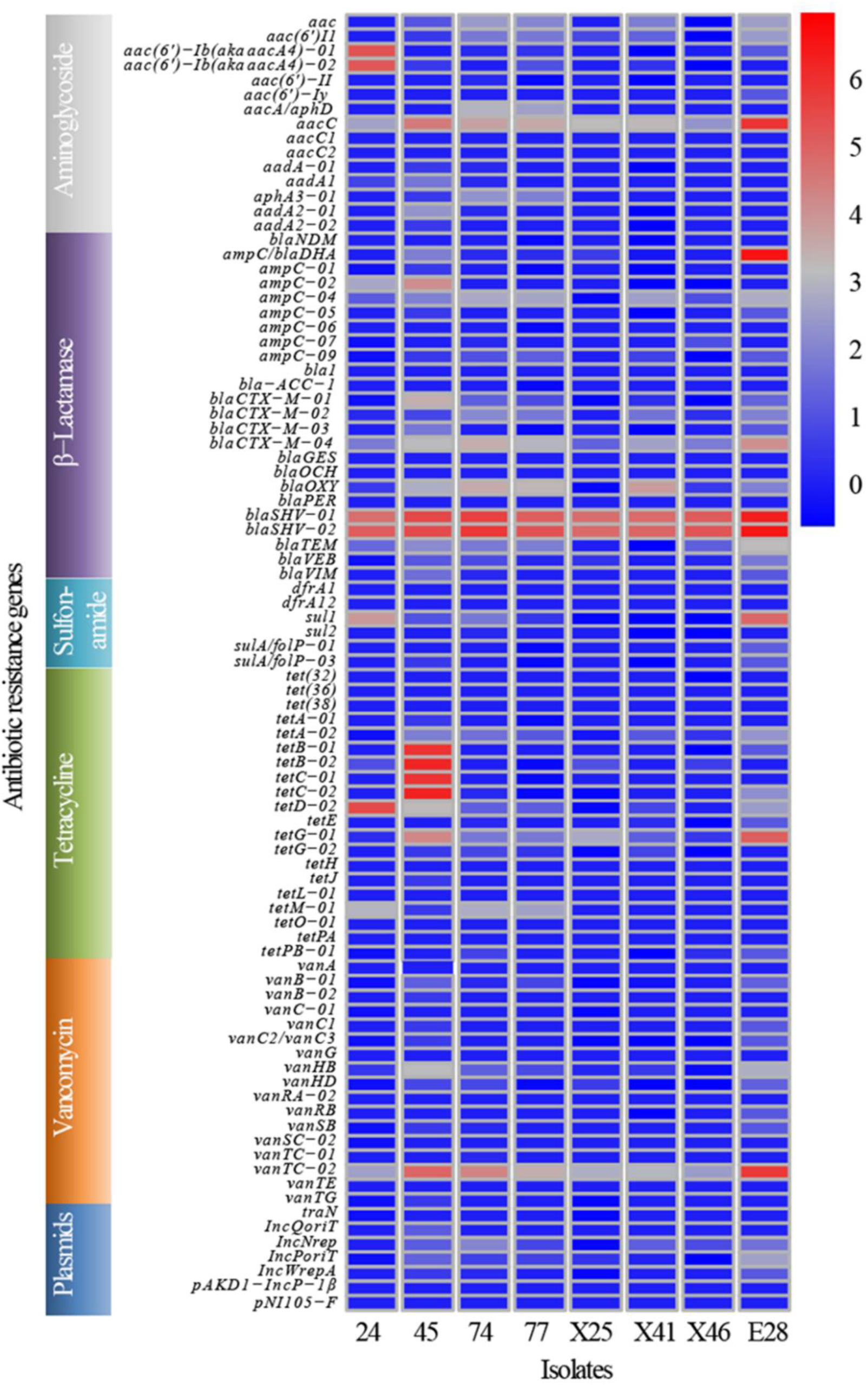
ARGs distribution in CRKP in giant panda. Columns: the number of isolates in the study; Rows: 89 different types of ARGs.

### 3.3 The virulence genes and MGEs of 8 CRKP isolates

The identification of virulence-related factors is a crucial step in comprehending the molecular basis of bacterial disease. Hence, 16 virulence genes from 8 isolates of CRKP were detected and analyzed in this study. Among them, 7 genes tested positive, including wabG (100.0%), uge (100.0%), ycf (100.0%), entB (87.5%), kpn (50.0%), alls (25.0%), and wcaG (25.0%). In addition, strain E28 exhibited the highest number of virulence genes with a total of 6 virulence genes present (wagGI, uge, alls, wcaG, kpn, ycf), while other strains carried only 4 or 5 virulence genes (Fig. 3).

**Fig. 3.**
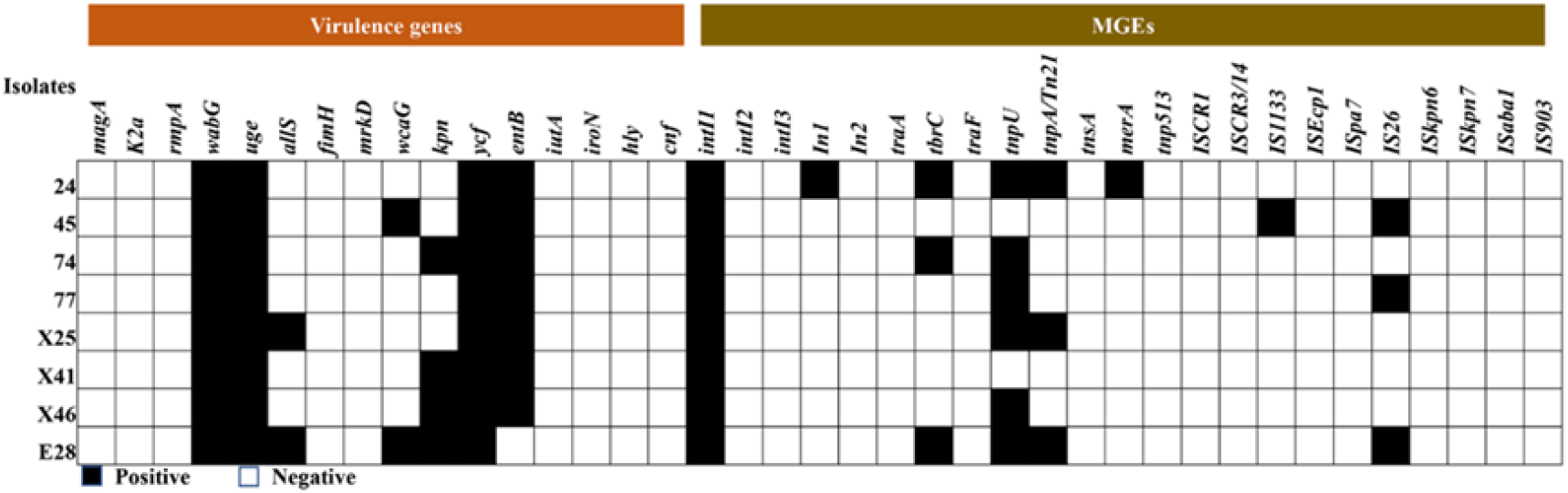
The detected result of virulence genes and MGEs in CRKP strains isolated from giant pandas. Columns: 16 different types of virulence genes and 23 different types of MGEs; Rows: the number of isolates in the study.

Besides,this study also included the analysis of a total 23 MGEs. The intI1 gene was detected in all 8 strains, while there were significant variations in the detection rates among the other 22 MGEs. Specifically, the detection rates of In1, tbrC, tnpU, tnpA/Tn21, merA, IS1133 and IS26 were found to be 12.5%, 37.5%, 75.0%, 37.5%, 12.5%, 12.5% and 37.5% respectively. None of the remaining MGEs were detected. Furthermore, there were notable differences in the number of MGEs carried by different strains, among which strain 24 carried the most MGEs species (6/23) (Fig. 3).

### 3.4 The molecular epidemiology of 8 CRKP isolates

The 8 CRKP isolates exhibited distinct PFGE patterns and highly diverse MLST types, as depicted in Fig. 4. These CRKP strains displayed a high level of genetic diversity, with less than or equal to 84%. MLST analysis revealed that 8 CRKP strains belonged to different sequence types (ST).

**Fig. 4.**
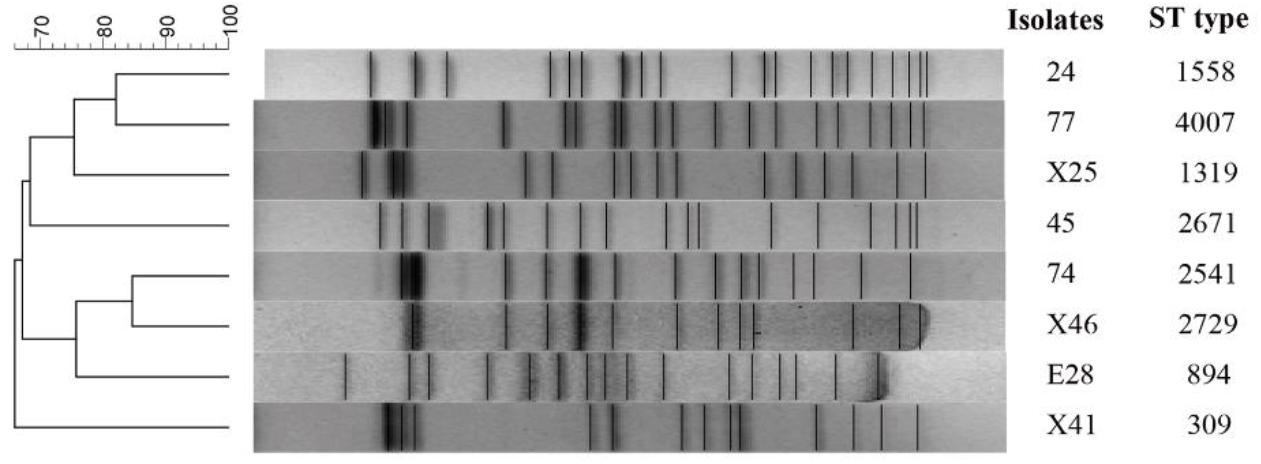
Molecular epidemiology of SRKP through the analysis of PFGE and MLST1. The dendrogram of PFGE, generated using the BioNumerics software, illustrated the relatedness of fingerprints among eight CRKP strains isolated from giant pandas. The dendrogram was constructed based on the restriction patterns of XbaI-digested KP genomes, with strain numbers and ST types depicted alongside the corresponding restriction profiles.

## 4. Discussion

The emergence of CRKP was initially documented in the United States, followed by subsequent reports from The French Republic, Sweden, and Canada (White, McIver et al. 2001, Chang, Sharma et al. 2021). In a hospital located in eastern China, the isolation rate of CRKP reached 41.3%, surpassing isolation rate 4.5% in our findings in giant pandas (Lu, Zhang et al. 2023). This disparity could potentially be attributed to the comparatively lower utilization of antibiotics for treating giant pandas, and the fact that our study isolated CRKP strains exclusively from fecal samples obtained from healthy giant pandas as opposed to clinical specimens collected in other investigations.

The antibiotic susceptibility test of CRKP isolates revealed that 87.5% of the isolates exhibited resistance to imipenem; while 25.0% showed resistance to doxycycline. Additionally, 12.5% displayed resistance to meropenem, cefazolin and chloramphenicol, indicating varying degrees of drug resistance among the eight CRKP isolates in this study, particularly towards carbapenems. Notably, the resistance of CRKP isolates to imipenem and meropenem varied across different countries and regions; for instance, in Saudi Arabia, the rates were 55.6% and 61.7%, respectively; whereas in New York, USA, they were observed at 17.0% and 20.0%; similarly in Chongqing, China, they stood at 37.2% and 30.8%, respectively(White, McIver et al. 2001, Chang, Sharma et al. 2021, Bshabshe et al. 2020; Kaiser et al. 2013). The resistance rate of the eight CRKP strains to imipenem in this study (87.5%) was significantly higher compared to that of meropenem (12.5%). The reasons for this outcome could be attributed to several factors: firstly, the utilization of antibiotics varied across different countries and regions, and the results of resistance test suggested that veterinarians may preferentially administer imipenem during clinical treatment of giant pandas; secondly, there existed disparities in the resistance mechanisms between the two CRKP strains, as meropenem targets both PBP2 and PBP3, whereas imipenem exhibits a stronger affinity towards PBP2 only (Zhanel et al. 2007). Additionally, it was worth noting that all eight CRKP strains examined in this study demonstrated sensitivity towards amitrazan, kanamycin, gentamicin, ofloxacin and norfloxacin. These findings suggested potential clinical options for preventing and treating bacterial infections caused by these strains.

HT-qPCR technique enables rapid and sensitive quantification of numerous ARGs that confer resistance to nearly all major classes of antibiotics and provide a comprehensive profile of ARGs (Su et al. 2015). Therefore, this method was employed to perform a total of 90 ARGs on 8 CRKP strains. Out of these, 47 ARGs were detected, with 6 ARGs positively presented in all strains, namely aacC, blaCTX-M-04, blaOXY, blaSHV-01, blaSHV-02 and vanTC-02. The most abundance ARGs identified included ampC/blaDHA, blaSHV-01, blaSHV-02, tetB-01, tetB-02, tetC-01, tetC-02, which was consistent with the findings reported by Hu (Hu et al. 2021). Simultaneously, the results from antimicrobial susceptibility testing revealed that the MDR isolates also exhibited significant resistance towards β-lactam antibiotics, particularly imipenem had an alarming resistance rate of 87.5%. The primary mechanism underlying KP’s β-lactam antibiotic resistance lies in its production of extended-spectrum β-lactamases encoded by extended-spectrum β-lactamase genes that disrupt the β-lactam ring structure leading to antibiotic inactivation (Fluit et al. 2001). The resistance phenotype and genotype of the isolates in this study closely corresponded, despite the diversity observed in both β-lactam antibiotics and genes. Given the rapid dissemination and emergence of novel ARGs among bacteria populations worldwide, it became imperative to monitor closely for any resistance genes carried by giant panda-associated bacteria.

MGEs play a crucial role in facilitating the transmission of ARGs within KP strains. In this study, a total of 23 MGEs were analyzed. It was observed that intI1 was present in all 8 strains, while there were significant variations in the detection rates of the other 22 MGEs. The detection rates for In1, tbrC, tnpU, tnpA/Tn21, merA, IS1133, and IS26 were found to be 12.5%, 37.5%, 75%, 37.5%, 12.5%, 12.5%, and 37.5%, respectively; whereas the remaining MGEs were not detected. Interestingly, different strains exhibited varying numbers of MGE species, with strain 24 carrying the highest diversity of MGEs. The emergence of multidrug resistance in microorganisms is believed to be closely linked to MGEs due to their ability to swiftly transfer multiple ARGs (Martinez, 2009). The horizontal transfer of ARGs mediated by MGEs is considered as a primary mechanism driving the spread of ARGs (Chen et al. 2018). MGEs, serving as carriers of ARGs, play a pivotal role in capturing, accumulating, and disseminating these genes. This transfer can occur both intra-strain and inter-strain within KP, thereby facilitating the rapid and widespread proliferation of antibiotic-resistant bacteria. Moreover, the horizontal transfer of ARGs contributes to the emergence of drug-resistant and multi-drug-resistant bacteria in clinical settings (Chamosa et al. 2017). In this study, we found that multidrug-resistant KP strains isolated from different giant pandas carried common ARGs along with a substantial number of MGEs detected. Therefore, it is plausible to hypothesize that there might be sharing of ARGs among the giant pandas in this study. However, further investigations were required to determine the precise mechanisms underlying the sharing of ARGs.

The detection of 16 common virulence genes from 8 CRKP strains was also an important part of this study. Seven genes, namely wabG (100%), uge (100%), ycf (100%), entB (87.5%), kpn (50%), all (25%), and wcaG (25%), were found to be positive. Additionally, among the strains, E28 carried the highest number of virulence genes, with 6 virulence genes identified as wagGI, uge, all, wcaG, kpn, ycf, while other strains carried only 4 or 5 virulence genes, indicating a relatively high prevalence of multiple-virulence-gene-carrying strains in adult giant panda-sourced KP. Genes associated with lipopolysaccharide (LPS) production primarily included uge and wabG, encoding UDP-galactose-4-epimerase and galactosyltransferase, respectively. These two types of virulence genes are commonly presented in a variety of virulence factors within KP strains. Studies have demonstrated that KP lacking the uge gene produces incomplete lipopolysaccharides (Regué et al. 2004). Mutations or deletions of the wabG gene in KP can result in the absence of certain capsular antigens and hemolysins, which reduces pathogenicity in animal infection models (Cogen et al. 2009). In addition, there are several other genes potentially associated with the virulence of KP, including kpn gene involved in capsule polysaccharide synthesis, and wcaG and wagGI genes encoding extracellular toxins. Research has suggested that genes related to iron carriers, pili, and lipopolysaccharides are located on virulence plasmids in highly virulent KP strains and serving as specific molecular markers for high-virulent strains (Russo et al. 2019; Xu Shuibao et al. 2017). Therefore, it can be inferred that wabG, uge, and ycf may potentially serve as high virulence genes. In this study, the LPS carrier genes wabG and uge exhibited a detection rate of 100%, which aligned with the findings of Zhang Wenju (Zhang Wenju et al. 2020). Conversely, unlike Han Kun’s study, the fimH pilus virulence gene was not detected in KP strains from different animal sources, indicating variations in the carried virulence genes among KP isolates. The iron carrier gene entB displayed a detection rate of 87.5%, while all urease gene had a detection rate of 25%. These results were consistent with those reported by Wang Zhehong (Wang Zhehong et al. 2021). This study identified one strain of isolated bacteria carrying 6 virulence genes, including lipopolysaccharide carrier genes and urease genes, suggesting its potential as a highly virulent KP strain isolated from giant pandas, laying the foundation for future investigations on the pathogenicity.

The DNA fingerprinting by PFGE is considered to be the “gold standard” for typing of microorganisms due to the adequate consistency within a single assay, which is widely used in molecular epidemiology (Jouni Heikkinen et al. 2022). MLST technique is a well-established procedure employed for characterizing bacterial species by sequencing internal fragments of typically seven housekeeping genes (Jolley et al. 2018). Eight strains of bacteria exhibited distinct PFGE patterns, indicating a high degree of genetic diversity (≤ 84%) among these CRKP strains. Further analysis using MLST revealed that the eight CRKP strains could be classified into different STs. This study successfully divided the eight CRKP strains into eight distinct STs, indicating that despite the observed diversity among CRKP strains isolated from various giant pandas, some strains still share a common origin.

## 5. Conclusion

In this study, we investigated eight strains of CRKP isolated from giant pandas to determine their antibiotic susceptibility, ARGs, MGEs, virulence genes, and molecular epidemiology. The most abundant ARGs included ampC/blaDHA, blaSHV-01, blaSHV-02, tetB-01, tetB-02, tetC-01, and tetC-02. Analysis of MGE revealed the presence of intI1 in all strains, while other MGEs exhibited varying detection rates. Strain 24 carried the highest diversity of MGE species. Seven virulence genes including wabG, uge, ycf, entB, kpn, alls, and wcaG were detected across the strains with varying proportions. Molecular epidemiology analysis using PFGE patterns indicated a high level of genetic diversity among the CRKP strains. MLST analysis classified the strains into different STs.

In conclusion, this study highlighted the remarkable diversity of CRKP strains in giant pandas, exhibiting varying degrees of antibiotic resistance and the presence of multiple ARGs and virulence genes. These findings emphasized the critical significance of monitoring and researching antibiotic resistance in wildlife populations to safeguard the well-being of these endangered animals.

## 6. ETHICS STATEMENT

The animal study was reviewed and approved by the Institutional Animal Care and Use Committee (IACUC) of the Chengdu Research Base of Giant Panda Breeding (No. 2018017).

## 7. AUTHOR CONTRIBUTIONS

Xia Yan, Xiaoyan Su, Rong Hou, Lin Li, Chanjuan Yue, and Songrui Liu contributed to conception and design of the study. Xiaoyan Su played a guiding role in carrying out the experiment. Xia Yan, Mei Yang, Yunli Li, and Dongsheng Zhang collected samples. Xia Yan, Mei Yang, and Yang Zhou performed bacterial identification and isolation and related component testing.Xiaoyan Su, and James Edward Ayala performed the statistical analysis. Xia Yan, Mei Yang, and Xiaoyan Su wrote the first draft of the manuscript. All authors contributed to manuscript revision, read, and approved the submitted version.

## 9. ACKNOWLEDGMENTS

We would like to thank the veterinary staff and keepers of the Chengdu Panda Base for collecting samples, and Cen Xin for arrangement of article data.

---

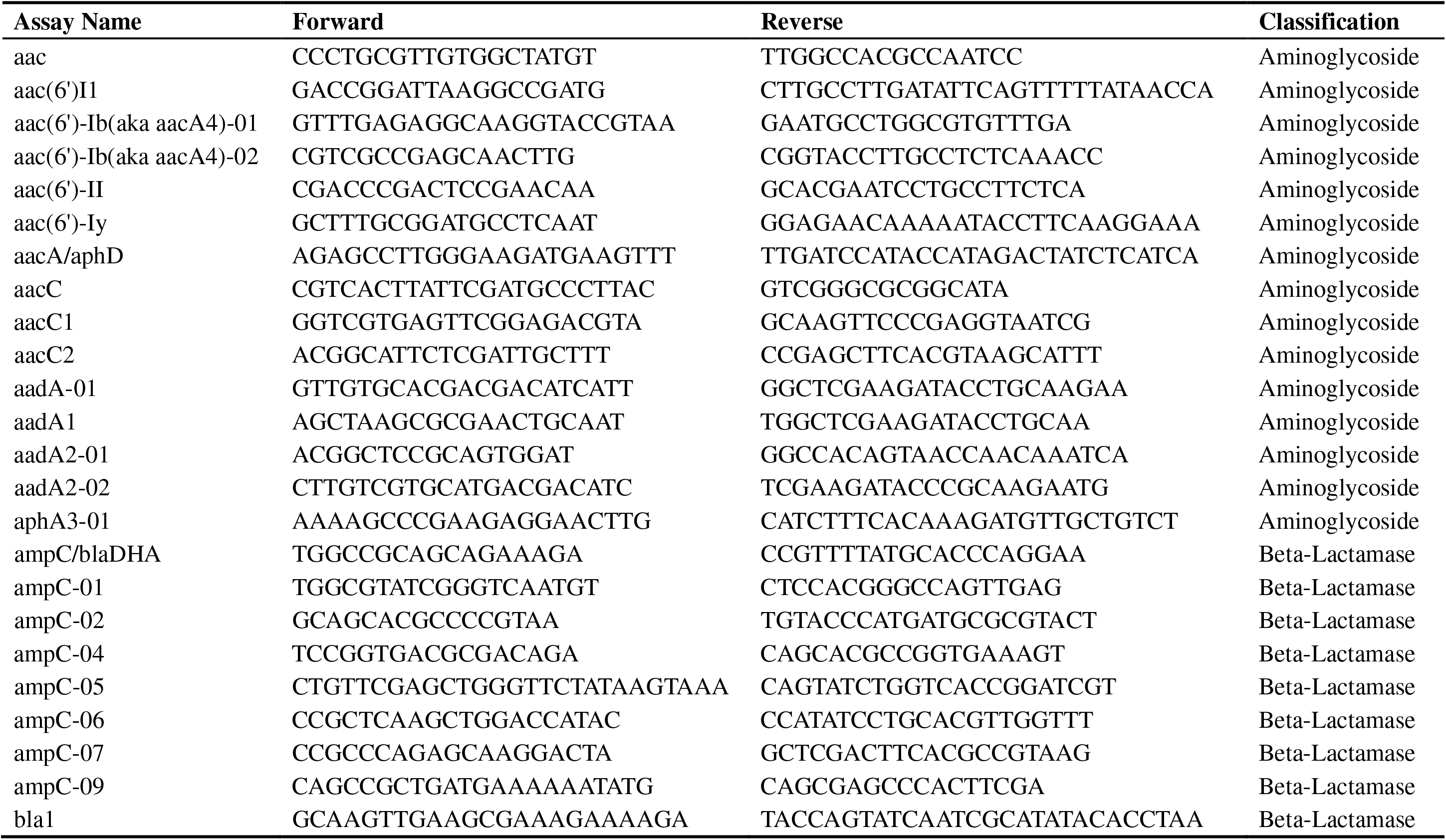
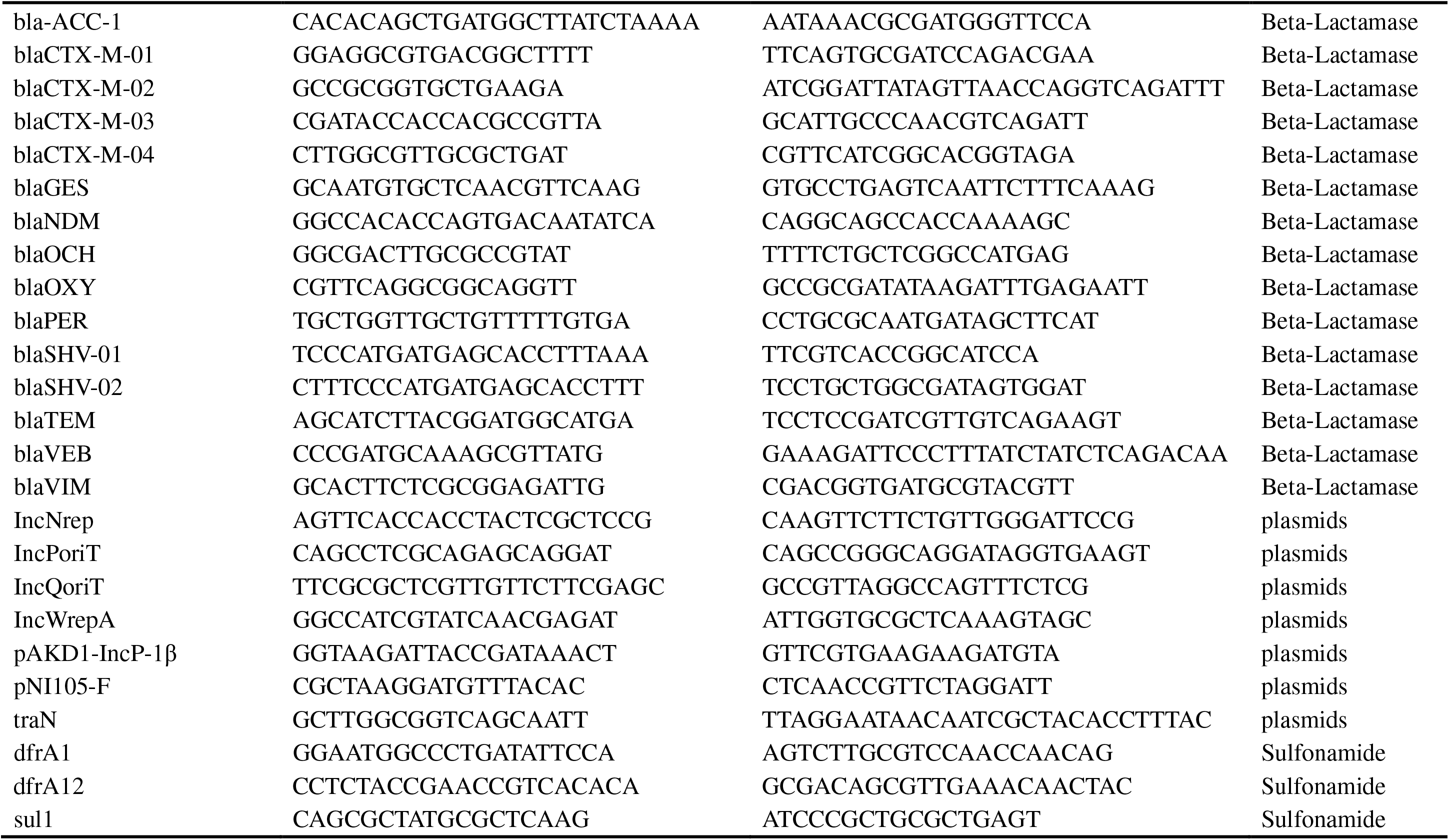
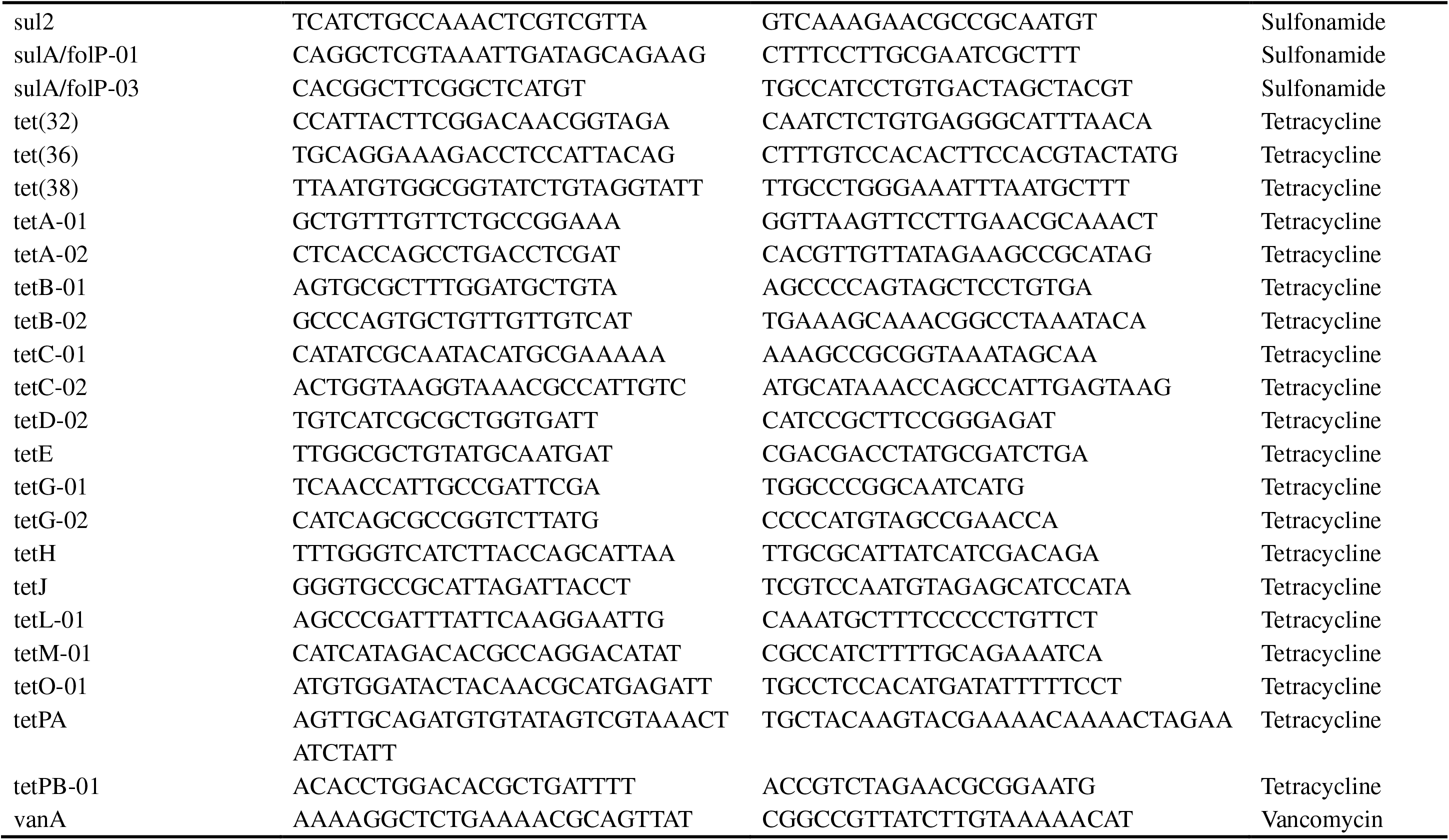
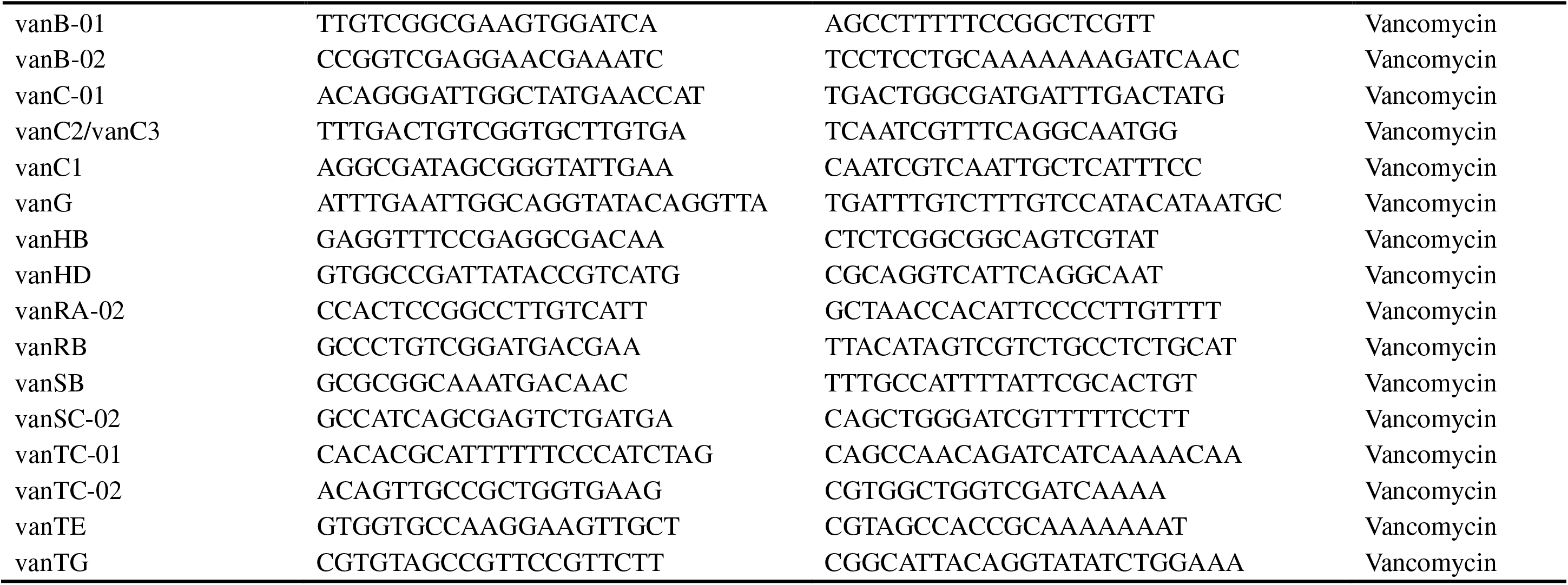

